# Presynaptic Actin Nanostructures: A Reproducibility Case Study

**DOI:** 10.64898/2025.12.18.695210

**Authors:** Rensu P. Theart, Florian Levet, Paul Hernández-Herrera, Christophe Leterrier, Laura R. de la Ballina

**Affiliations:** Department of Electrical and Electronic Engineering, Stellenbosch University, Cnr Banghoek Road & Joubert Street, Stellenbosch, 7600, South Africa; Univ. Bordeaux, CNRS, Interdisciplinary Institute for Neuroscience, IINS, UMR 5297, F-33000 Bordeaux, France; Univ. Bordeaux, CNRS, INSERM, Bordeaux Imaging Center, BIC, UAR3420, US 4, F-33000 Bordeaux, France; Universidad Autónoma de San Luis Potosí, Facultad de Ciencias, San Luis Potosí, SLP, México; Aix Marseille Université, CNRS, Institut de neurophysiopathologie, UMR7051, NeuroCyto, Marseille, France; Centre for Cancer Cell Reprogramming, Institute of Clinical Medicine, Faculty of Medicine, University of Oslo, Montebello, 0379, Oslo, Norway; Department of Molecular Cell Biology, Institute for Cancer Research, The Norwegian Radium Hospital, Oslo University Hospital, Montebello, 0379, Oslo, Norway

**Keywords:** bioimage analysis, reproducibility, GloBIAS, actin nanostructures, presynapses

## Abstract

In an effort to assess the reproducibility of bioimage analyses in current publications, we took part in a Global BioImage Analysts’ Society (GloBIAS) initiative to try and reproduce results from published articles. We attempted to reproduce core findings from the work of Bingham *et al.*, which investigates the actin organisation in presynaptic structures by using diffraction-limited and super-resolution microscopy. While the original paper unveiled clear biological insight, it lacked sufficient detail in the bioimage analysis methodological approach, limiting the depth of reproducibility we could achieve. Through frequent contacts with the corresponding author, we managed to replicate qualitative aspects of the analysis of actin nanostructures in bead-induced presynapses. We performed image reconstruction from super-resolution microscopy data, automatic image registration and visual inspection, followed by manual annotation of structures of interest in ∼35 images. Our experience with this exercise highlights the importance of transparent data sharing and accessibility, as well as the need to adhere to bioimage analysis standards that ensure the reproducibility of nowadays complex biological image analysis studies. It also shows how crucial interdisciplinary collaboration is, since many biology labs are simply unaware of these standards, which are often easy to implement and would likely be widely adopted if their value was better understood or known.

## INTRODUCTION

As part of the Global BioImage Analysts’ Society (GloBIAS) first in-person workshop (*GloBIAS Bioimage Analysts Workshop*, November 2024, Gothenburg, Sweden), we conducted a practical exercise addressing “Bioimage analysis workflows”, in which workshop participants were divided into working groups and assigned published articles where bioimage analysis workflows had been used. The assignment consisted of trying to reproduce, based on the *Methods* described in the paper, the published bioimage analysis workflows and presenting the results during a dedicated session in the GloBIAS workshop, with the final aim of assessing how reproducible bioimage analysis workflows are at present.

In recent years the bioimage analyst community has generated guidelines to follow when publishing bioimages and bioimage analysis workflows, in hope of increasing the rigor and reproducibility of published images and corresponding extracted data and with the aim of producing scientific data more aligned with the FAIR principles^1–3^. We used the “*Checklists for publication of image-analysis workflows*” included in those guidelines to assess the reproducibility of our assigned article^1^.

After two failed attempts (due to lack of access to original data) with other articles, we ended up assessing the reproducibility of the work published by Bingham *et al.* which revolves around understanding the organisation of actin nanostructures in presynapses^4^. In this paper, the authors exploit a previously validated model where presynapses are induced in cultured neurons using polylysine-coated beads^5^. By studying the induced presynaptic structures using Single Molecule Localisation Microscopy (SMLM), the authors identify three distinct actin nanostructures in presynapses: actin mesh, actin rails and actin corrals, which they further confirm to be present in natural presynapses.

From this extensive work (the original article contains 10 main figures and 7 supplementary ones), we selected *Figure 7* as the subject of our reproducibility exercise. In that figure, the authors confirm the presence of *actin mesh, rails,* and *corrals* in various types of induced presynapses. Our task was to assess whether – based on the analysis steps and textual description published by the authors– we were able to identify the same structures they did.

After getting in contact with the corresponding author of the original publication and having access to sample image data and a more detailed description of the actin nanostructures of interest, we proceeded with our annotation exercise.

We were partially successful in following the author’s instructions and in identifying actin nanostructures in bead-induced presynapses, with discrepancies in the results likely due to the different backgrounds of each team member.

During this reproducibility exercise, we uncovered several limitations in the reproducibility of bioimage analysis workflows, many of which could be avoided by adhering to the standards and guidelines of the bioimage analysis community.

## METHODS

### Contacting the corresponding authors to gain access to the original data

For all articles considered in this reproducibility assignment, the lack of directly available image and/or code data required contacting the corresponding authors of the original works (as stated in the “*Data and Code Availability*” sections of the original publications) to obtain a dataset we could work with.

In the case of the work by Bingham *et al.*, after contacting the corresponding author we gained access to sample SMLM image data – 34 regions of interest (ROIs) from 3 replicates (NC1, NC2 and NC3) – and the authors’ annotations for those same ROIs (which we only consulted to contrast results once we had blindly performed our analysis) first and the corresponding epifluorescence images later on (as we realised we were missing some relevant information we could not extract from only the SMLM images). We have deposited those images in Zenodo and are now publicly available (at https://doi.org/10.5281/zenodo.17792816)^6, 7^.

We needed several rounds of communication with the corresponding author to get clarifications regarding the analysis steps, as well as the definitions of the different presynaptic actin nanostructures we needed to identify.

Due to the essential role of the corresponding author of the original publication in enabling the reproducibility of the bioimage analysis workflow evaluated, they are included in this work’s author list. We wish to state that this inclusion did not affect our assessment of the reproducibility of the original work.

### SMLM image reconstruction

Following the indications in the ChriSTORM (version 1.3) GitHub repository (https://github.com/cleterrier/ChriSTORM) referred to in the original work of Bingham *et al.*, we used the version of the ThunderSTORM plug-in for ImageJ by Koen J.A. Martens (which can be found at https://github.com/kjamartens/thunderstorm) to process data acquired with single-molecule localisation microscopy (SMLM) (available at https://doi.org/10.5281/zenodo.17792816, under *00_raw_images*, contained in *Locs_TS_ROIs* folders)^8, 9, 7^. ChriSTORM provides macros for converting, filtering, and batch-processing localisation data and for generating reconstructed images, thereby complementing the ThunderSTORM plugin.

In FIJI (using ImageJ2, version 2.14.0/1.54f) we selected “Plugins > ThunderSTORM > Import/Export > Import results” to import the comma-separated values (.csv) files from the “*Locs_TS_ROIs*” folder of each replicate^10, 11^. In the “Camera setup” menu, we adjusted the “Pixel size [nm]” to 160 and pressed OK. In the “Input File” section, we selected *CSV (comma separated)* as “File format” and in “File path” we indicated the path to the “*Locs_TS_ROIs*” folder containing the .csv files (folder browsing is accessible after clicking the “…” button) and pressed OK. In the “ThunderSTORM:results” window, we clicked the “Visualization” button, where we selected *Auto size by results* in the “ROI” section, and under the “Visualization options” section, we selected *Gaussian reconstruction* as Method. After unticking the “3D” checkbox, we clicked OK to get the SMLM reconstructed image (**Figure 1**).

**Figure 1.**
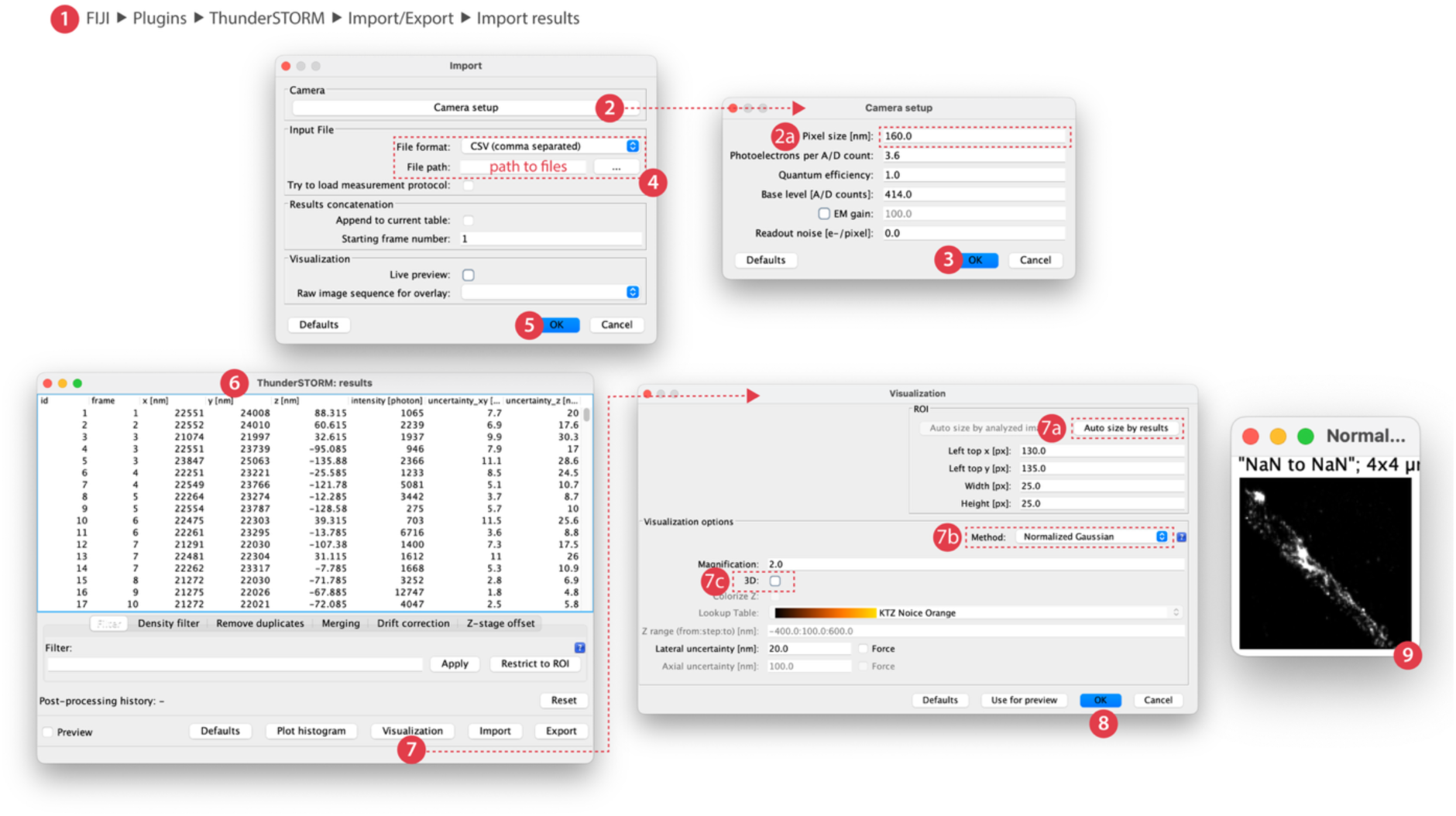
SMLM image reconstruction using the ThunderSTORM FIJI plugin. Sequential steps to use the ThunderSTORM FIJI plugin to reconstruct single molecule localisation microscopy images. After starting the plugin (1), we access the “Camera setup” menu (2) to adjust the pixel size (in nm) (2a) and press OK (3). We then proceed to adjust the “Input file” section, selecting *CSV (comma separated)* as File format and indicating the local path to the corresponding .csv files (4). Once this is in place we click OK (5) and are presented with the ThunderSTORM:results menu (6). We click “Visualization” (7) and in the appearing window we click “Auto size by results” (7a) choose *Normalized Gaussian* method (7b) and untick the 3D box (7c). After clicking OK (8) we get the SMLM reconstructed image (9). In the figure we present screenshots from FIJI.

These steps produced super-resolution renderings of actin localisation patterns from the raw SMLM data (included in https://doi.org/10.5281/zenodo.17792816, under *01_processed_images* and *Recs_TS_ROIs* folders, for each replicate) ^7^. However, ChriSTORM and ThunderSTORM are limited to data processing and visualisation; they do not include tools for ROI definition, multimodal image registration, or structural classification. As such, the shared code facilitated reproducible image reconstruction but did not encompass the analytical steps necessary to identify or quantify specific presynaptic actin nanostructures described in the original study.

### Description of our structures of interest: presynaptic actin nanostructures

Our aim was to annotate the ROIs we had been provided for the corresponding structures of interest. It is worth noting that in the original publication these structures had been defined following an unbiased consensus process, where two specialists independently annotated ∼50 synapses, and then the authors defined as *typical* structures the ones that were frequently observed by both specialists. Thus, the criteria defined by the authors to identify the three types of presynaptic actin nanostructures were as follows: a) *actin mesh*: a small cluster of actin opposed to the bead contact and within the presynaptic marker cluster, b) *actin rails*: linear structures within the presynaptic marker cluster, *actin corrals*: large actin clusters at the periphery or just outside of the presynaptic marker cluster.

Due to our lack of experience in the field, we had to confirm with the authors that the “presynaptic marker cluster” corresponded to the synaptophysin label. We realised that we were missing that data, as well as information related to “bead contact”. We got back in contact with the authors, asking for extra sample image data we could extract that information from and were provided with fluorescent images for each replicate (which can be found at https://doi.org/10.5281/zenodo.17792816, under *00_raw_images* and each corresponding replicate folder: *NC1_*, *NC2_* and *NC3_epifluo.tiff* files)^7^.

### Automatic registration of epifluorescence and SMLM images and generation of multimodal pseudo-image

After obtaining the epifluorescence images corresponding to the ROIs where they had performed SMLM acquisitions (actin STORM images), we could get information related to presynaptic marker cluster synaptophysin (channel 3 in epifluorescence images), axon marker ß2-spectrin (channel 4 in epifluorescence images) and the presynaptic-inducing beads (channel 6 in epifluorescence images.

To segment both synaptophysin and bead markers, we used wavelet analysis and WaveJ (https://github.com/flevet/WaveJ), an update of the wavelet component of SpineJ (https://github.com/flevet/SpineJ) that was extended to handle image stacks^12^. After adding the *wavej_package.jar* file –which can be found as part of WaveJ release 1.0.0 (https://github.com/flevet/WaveJ/releases/tag/1.0.0)– to your computer’s FIJI “*plugins*” folder, WaveJ will be available as a FIJI plugin. Open the image (individual channel) containing the objects you want to segment and run WaveJ. The plugin uses “à-trous” wavelet algorithm, which computes a series of multi-scale wavelet coefficients by iterative convolutions of increasing kernels^13^. It is then sufficient to filter those coefficients at the level corresponding to the size of the objects of interest to reliably segment them. For the synaptophysin segmentation, we used the wavelet coefficient level 1 with a threshold of 5 and a minimum size for the segmented clusters of 10 pixels. For the bead segmentation, we used the wavelet coefficient level 2. This time we had to slightly modify the thresholds used depending on the signal to noise ratio and image quality of the different replicates. For thresholding the coefficients, we used two values: 10 for NC1, NC2 and NC3 images 1 to 8, and 3 for NC3 images 9 to 15. For the minimum size of the beads, we also used two values: 80 for NC1 and 60 for NC2 and NC3 (all images) (**Figure 2** exemplifies the steps followed to achieve bead segmentation on channel 6 of *NC1_epifluo.tiff*).

**Figure 2.**
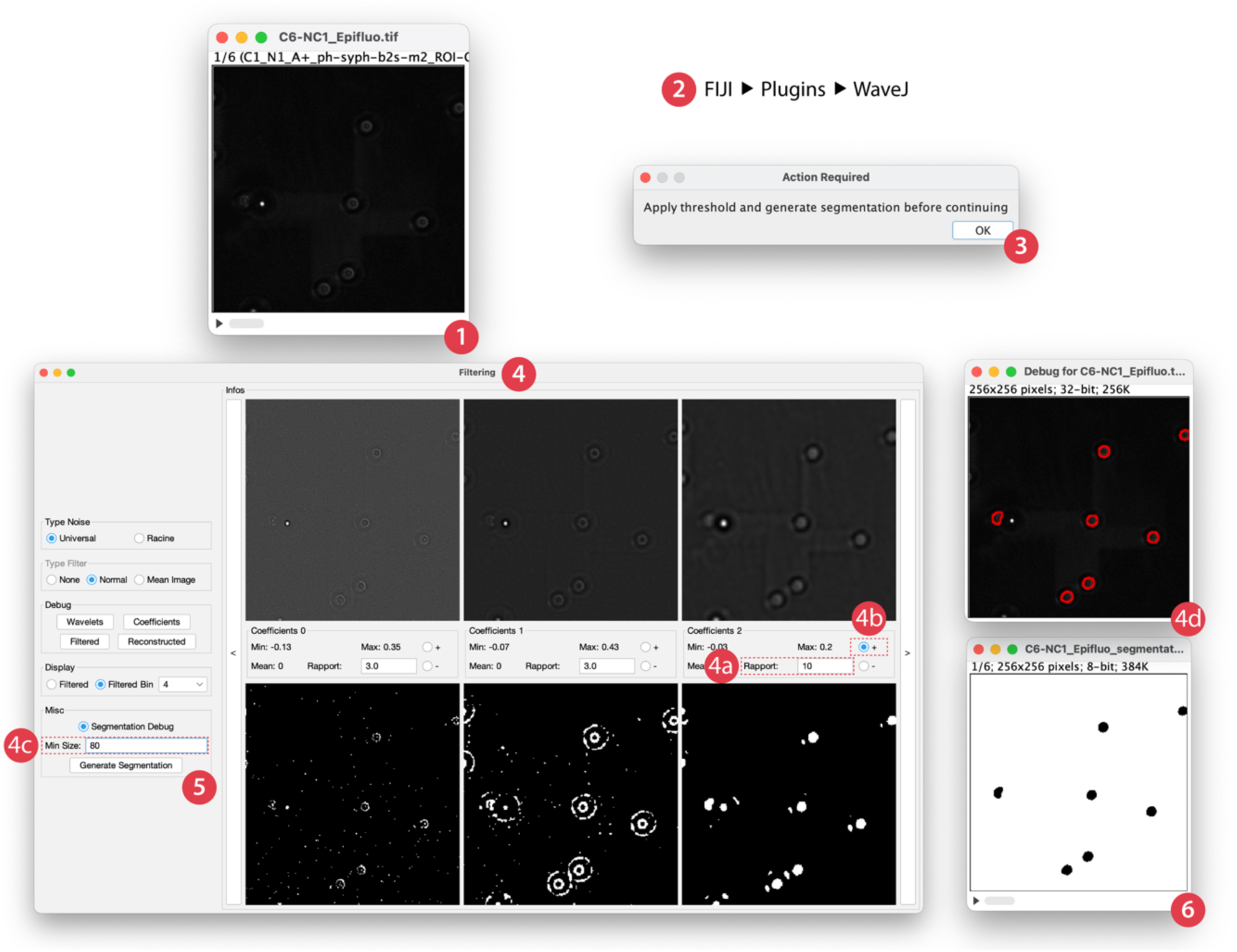
Epifluorescence image segmentation using the WaveJ FIJI plugin. We start from the individual fluorescent channel we want to segment (channel 6, corresponding to beads in this case)(1) and open the WaveJ FIJI plugin (2). We get a dialog asking to apply a threshold and generate a segmentation, we can click OK (3) The WaveJ “Filtering” panel (4) displays wavelet coefficients at different scales. The bottom row shows the resulting binarized masks for each scale. After choosing a threshold factor for a specific coefficient (4a) we choose to add all white pixel of that wavelet level from the final image (4b). We need to input the minimum area (in pixel^2^) for the objects we want to segment, beads in this case (4c). The debug window shows and updated output representation as we are adjusting parameters (4d). Once we are done with the settings we click on “Generate Segmentation” (5) to create the final binarized image (6), which is ready for export and downstream comparison. In the figure we present screenshots from FIJI.

We then upsized by a factor of 4 the epifluorescence images and downsized by a factor of 10 times the SMLM ROIs, bringing both images to a 40 nm pixel size. We created a RGB stack (included in https://doi.org/10.5281/zenodo.17792816, under *01_processed_images* and each replicate folder: *NC1_*, *NC2_* and *NC3_bead_synaptophysin_segmented.tiff* files) where we combined the information from the actin SMLM acquisition (channel 1 in RGB stack), with the ß2-spectrin image from the epifluorescence stack (channel 2 in RGB stack), and the combined contours of beads (intensity 125) and synaptophysin (intensity 255) segmented from the epifluorescence images (channel 3 in the RGB stack) ^7^. This stack was automatically obtained by extracting the crop coordinates encoded in the SMLM filename, allowing precise alignment within the larger field of views of the 3 other channels.

### Annotation of structures of interest

Once we had the pseudo-images that combined information from SMLM and fluorescence microscopy, we were ready to identify the distinct presynaptic actin nanostructures the authors had described in their original publication (see definitions above).

We started with each analyst from our working group separately annotating each ROI with what we understood to be actin mesh, actin rails and actin corrals (annotations can be found at https://doi.org/10.5281/zenodo.17792816, under *02_annotated_images/02_analysts_independent_annotations/* where we have a folder per analyst including their individual manual annotation files per replicate and ROI). We then contrasted our results and reached a consensus annotation per ROI (at https://doi.org/10.5281/zenodo.17792816, under *02_annotated_images/01_analysts_consensus_annotations/* where per replicate annotations are stacked in *.tif* files; folders including individual manual annotation files per replicate and ROI are also provided). These consensus annotations were then compared to annotations for each ROI that the authors had shared with us (and that we had not previously examined to avoid bias when generating our annotations, and which can be found in https://doi.org/10.5281/zenodo.17792816, under *02_annotated_images/00_Bingham_original_annotations* as per replicate stacked *.tif* files, both as originally provided by the authors and adjusted to follow the same colour code than the analysts’ annotations –see **Figure 3A**– in order to facilitate comparisons) ^7^.

**Figure 3.**
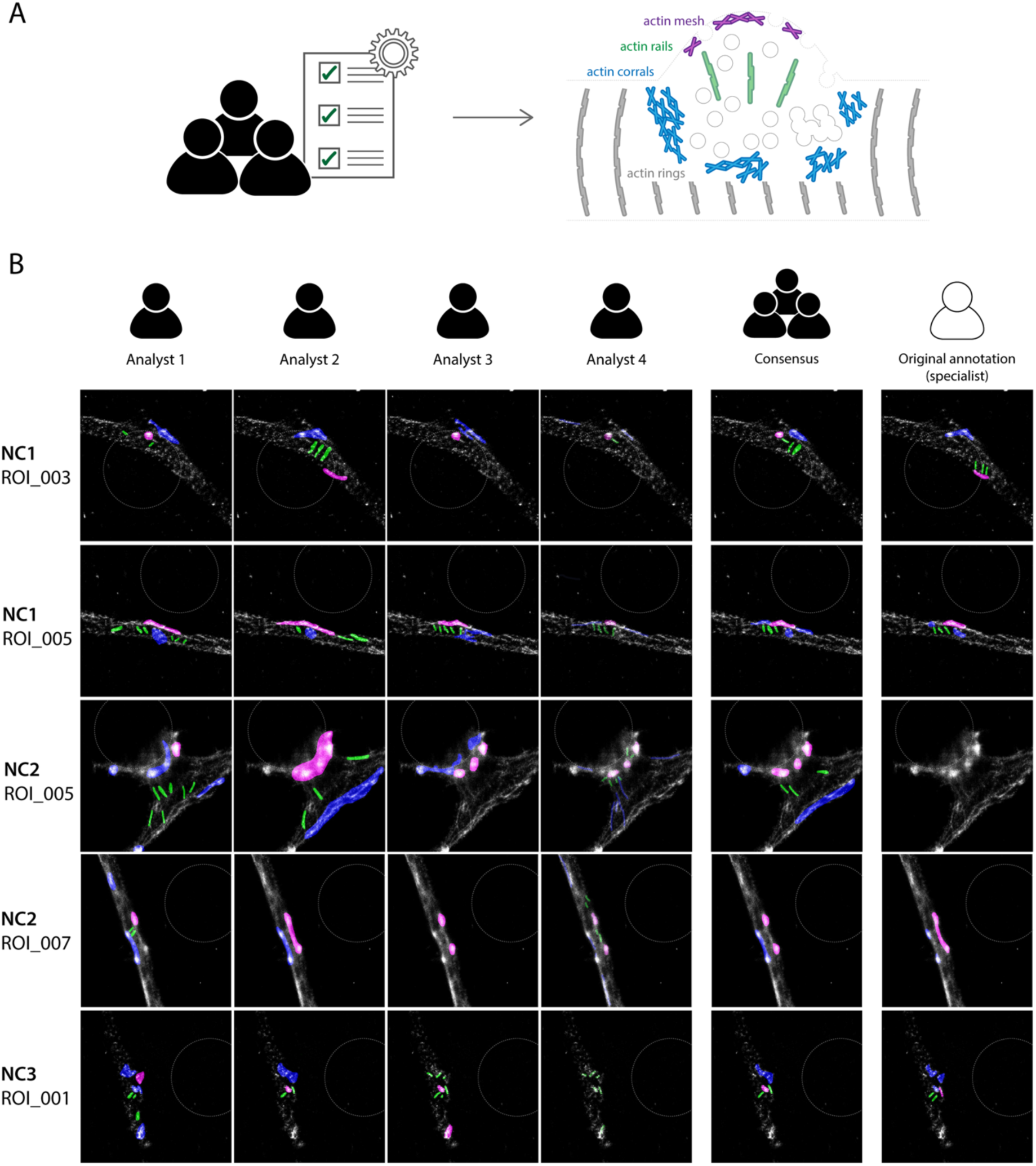
Actin nanostructure manual annotation. **A)** We aimed at identifying and annotating the different types of actin nanostructures (actin mesh in magenta, actin rails in green and actin corrals in blue) described in Bingham *et al.* by following the bioimage analysis workflow described in the Method section of the original publication. The right part of the panel has been adapted from *Figure 10G* from the original publication with kind permission from the authors^4^. **B)** Each of the analysts in this work performed independent annotations (colour codes as in panel A) for 34 ROIs (where bead position is indicated by a dotted circle). Individual annotations were compared and we agreed on a *Consensus* annotation for each ROI that was then contrasted with the *Original* annotations from the authors of Bingham *et al.* (specialists in synapse biology).

## RESULTS

The published work this exercise was performed on constituted the third attempt to find a publication with available sample image data and analysis code.

Our first assigned article did not provide access to any data or code; it had two sections specifically addressing *Data Availability* and *Code Availability*, which stated that both were available from the corresponding author upon reasonable request^14^. Unfortunately, our attempts to contact the corresponding author yielded no results and we were therefore forced to move to another article. The second work we were assigned included source code for image analysis available as a GitHub repository (https://github.com/JakubCzuchnowski/Supporting-data-for-Sensing-their-plasma-membrane-curvature-allows-migrating-cells-to-circumvent-o) but unfortunately, sample image data were not accessible^15^. We contacted the corresponding author, as stated in the *Data Availability* section, and this time we got a swift reply. Despite the authors’ willingness to provide original data, several constraints — the corresponding author was attending meetings and would be away for several days, the first author had moved overseas and had no access to the original data, at the host institution the raw data (large files) were backed up in tape and it would require some time to retrieve them — made us move to another article to ensure that we could complete the assignment in time for the *GloBIAS Bioimage Analysts Workshop*.

As in the previous cases, the work from Bingham *et al.* provided no access to sample image data we could analyse, although some code was accessible through a GitHub repository (https://github.com/cleterrier/ChriSTORM) ^4^. This code, however, only enabled reproducing the STORM image reconstruction, but did not include downstream annotation or ROI-based classification. Once again, the article’s *Data availability* section stated that data were available from the corresponding author upon reasonable request. This time around, after contacting the corresponding author, we got a prompt reply and gained access to original data to work on.

As a first approach to assess the bioimage workflow presented in the original work we contrasted it with the “Checklists for publication of image-analysis workflows”, which was published together with an online version (https://quarep-limi.github.io/WG12_checklists_for_image_publishing/analysis_workflows/intro_analysis_workflows.html) by members of Quality Assessment and Reproducibility for Instruments & Images in Light Microscopy (QUAREP-LiMi)^1^. Results of that assessment, where the original publication only scored 0.5 out of 9 points, are summarised in **Table 1**.

**Table 1.** *QUAREP-LiMi WG12 Checklist* applied to Bingham et al. (2023) “Presynapses contain distinct actin nanostructures” gives an overall assessment of 0.5/9, with scores of 0.5/5 associated to minimal requirements (1-5), and 0/2 for both recommended (6-7) and ideal (8-9) features.

| Checklist Item | What the Guideline Requires | How Bingham et al. (2023) Performed | Compliance Level (✓ / ⚠ / ✗) | Notes & Implications for Reproducibility |
| --- | --- | --- | --- | --- |
| **1. Cite workflow & platform** | Explicit citation of the full analysis workflow / software / environment. | Image reconstruction attributed to **ThunderSTORM** (Ovesný 2014) and “custom ImageJ macros (ChriSTORM)” by the corresponding author, but not cited as a citable software object or version. | ⚠ Partial |  |
| **2. Key settings** | Report variable parameters that affect outcomes (e.g., pixel size, thresholds). | No parameter values provided in the publication. | ✗ |  |
| **3. Example data** | Provide test dataset to validate workflow. | No sample data included; “Available upon request.” | ✗ | After request we managed to gain access to the raw data. |
| **4. Manual ROIs** | Include manually selected regions as separate files. | Manual annotations performed by authors but not deposited or referenced. | ✗ | After request we managed to get one example set of ROIs, but without explanation of why those ROIs were deemed valid. |
| **5. Exact version** | Report version of code or commercial software. | No version identifiers for ImageJ, ThunderSTORM, or ChriSTORM. | ✗ |  |
| **6. All settings (recommended)** | Provide or export full configuration/macro list. | Not supplied. | ✗ |  |
| **7. Public example (recommended)** | Make an example dataset publicly accessible. | None deposited. | ✗ | All replication attempts require personal communication. |
| **8. Document usage (ideal)** | Show screen captures / videos / tutorials. | None provided. | ✗ | Only some examples of the structures were given in the paper, but explanation of definition was not clear to non-biologists. |
| **9. Cloud container (ideal)** | Provide containerised / cloud-hosted reproducible environment. | None provided. | ✗ |  |

The segmentation of presynaptic actin nanostructures presented in *Figure 7* of the original publication had been done manually following visual inspection of the images^4^. In the *Methods* section, the authors provided a description of the criteria followed to identify the nanostructures, with some of the described features being difficult to identify for non-specialists in neuron biology. Given the limitation on the number of sample images presented in *Figure 7* and the lack of access to sample image data, we had to get in contact with the authors to get the raw image data first, and to clarify the criteria and steps followed to identify the actin nanostructures then.

Each analyst from the working group started by annotating images independently, and then results were put in common to reach a consensus annotation, which was contrasted with the annotations that the authors from the original work had shared with us (it is worth emphasising that, in order to avoid interference with our analysis, these validated annotations had not been consulted prior to this contrasting step).

When trying to reach a consensus annotation, we realised that there were many inconsistencies in the results obtained by the different analysts. We soon noticed that there were slight differences in the way each analyst interpreted the author’s description, which affected the results obtained when annotating the images (e.g. regarding the definition of *mesh* nanostructure: while one analyst interpreted it as “single brightest actin spot near bead”, another one considered it as “bright cluster inside the synaptophysin segmentation“).

When comparing our results with those from the original publication (**Figure 3** for representative examples and at https://doi.org/10.5281/zenodo.17792816, under *02_annotated_images/* where annotated stacked *.tif* files can be compared between *00_Bingham_original_annotations* and *01_analysts_consensus_annotations* folders for all ROIs) there were examples where there was a clear consensus among the analysts and the annotation nicely fitted with the one generated by the authors (**Figure 3**, NC1, ROI_005). In other cases, reaching a consensus annotation within the analyst group was more challenging, but in the end, the consensus was also in agreement with the authors’ annotations (**Figure 3**, NC3, ROI_001). Strikingly, there were cases in which the analysts easily agreed on a consensus that nevertheless entirely differed from the authors’ annotation (**Figure 3**, NC2, ROI_005). When we consulted the author’s notes, we realised that in some cases they had discarded specific ROIs for reasons like “the neuron did not look healthy” or “the synapse had an odd shape”, features that were not obvious to identify for the working group (composed primarily of image analysts and computer scientists) ^7^.

In all, our reproduction was qualitative and partially successful. We were able to identify nanostructures resembling the three categories described in the original work, and some of our annotations overlapped with those of the original authors.

## DISCUSSION

Despite the community recommendations of making data and code available through public repositories, during this exercise we experienced the challenges of getting access to original images and analysis pipelines^1^. Not having access to data makes it impossible to verify a pipeline’s robustness, and therefore we had to change twice the assigned article before gaining access to some original data we could work on.

In the case of the article by Bingham *et al.* no sample images that could be independently analysed were provided with the publication, which made difficult not only the attempt to reproduce the original bioimage analysis workflow, but also the publication of this article, as the images we wanted to annotate needed to be made publicly available (https://doi.org/10.5281/zenodo.17792816) before we could show the annotation results ^7^. The corresponding author of this publication was responsive and willing to help us gain access to all the information required to reproduce their analysis, noting that most of it had been performed manually. Whereas performing an automatic quantification would have obviously been better, this is something that cannot always be applied, especially when addressing challenging biological questions and using SMLM at the limit of its capacity.

In the case of this article, despite the availability of some code through a GitHub repository, steps followed to perform image registration, preprocessing, and annotation strategies were not described in sufficient detail, creating ambiguity and leading to variance in our reproduction efforts.

A big part of the original pipeline relied on manual inspection, which was difficult to replicate as the original work did not provide consistent criteria or enough training examples. As a result, there were annotation discrepancies among our working group members. While these differences may reflect subjective judgment, they also highlight the ambiguity in structure definitions—something that could be mitigated with more detailed and standardised methodological descriptions.

More broadly, defining robust and generalisable annotation guidelines is inherently challenging in biology due to the high variability of biological structures and imaging conditions. In many cases, large and diverse datasets are needed to capture this variability and establish clear criteria. In this study, the difficulty of acquiring the data limited the size of the available dataset, making it harder to establish reliable object definitions and contributing to differences in annotation outcomes.

As a result, the nanostructures identified in our reproducibility study were not always fully aligned with those described by the original authors. While part of these discrepancies may be attributed to our limited experience with neuronal data, it is worth noting that even domain experts often show variability when annotating complex biological structures^16^. In our case, this is further compounded by the fact that we were not specialists in synapse biology. For example, the original authors excluded some images from analysis based on subtle structural cues (such as assessing neuron health), which are difficult for non-experts to reliably evaluate.

Taken together, these observations underscore the importance of providing comprehensive annotation guidelines and considering inter-rater variability when manual inspection is part of the analytical process, especially in fields where expert interpretation plays a critical role.

It is worth mentioning that the original paper was written primarily from a biological perspective, with minimal emphasis on image analysis reproducibility. This highlights the importance of better communication between the bioimage analysis and biology communities, particularly regarding the existing guidelines that support reproducible practices. These guidelines are often straightforward to implement, and many biology labs would likely adopt them if they were more widely known and understood.

Despite the aforementioned limitations, the results from our exercise confirm that with clear methodological steps and defined structural definitions, some level of reproducibility is possible, even when assessing manual analyses. Our experience also highlights how collaborative discussion –especially with original authors–improves interpretation accuracy.

Ultimately, our bioimage analysis workflow reproducibility attempt confirms the importance of open access to sample image data, methods and code, as well as to intermediate analysis steps. It also underscores the need for dedicated bioimage analysts working alongside biologists from project inception to ensure that studies, including bioimage analysis workflows, comply with community standards and are rigorous and reproducible.

## CONFLICT OF INTEREST

The authors declare no conflict of interest.

## FUNDING

This project has been made possible in part by grant numbers 2023-321238 and 2023-329644 from the Chan Zuckerberg Initiative DAF, an advised fund of Silicon Valley Community Foundation. F.L. was supported by the Ministère de l’Enseignement Supérieur et de la Recherche (ANR-20-CE11-0006 NANO-SYNATLAS). C.L. was supported by Fondation pour la recherche médicale (FRM EQ20210301296). L.R.B was supported by the Research Council of Norway through FRIPRO grant number 314684.

## AUTHOR CONTRIBUTION

**Analysis**: RP Theart, F Levet, P Hernández-Herrera, LR de la Ballina; **Experiments**: C Leterrier; **Methodology**: RP Theart, F Levet, P Hernández-Herrera, LR de la Ballina; **Resources**: C Leterrier; **Software**: RP Theart, F Levet, P Hernández-Herrera, LR de la Ballina; **Supervision**: RP Theart, F Levet, P Hernández-Herrera, LR de la Ballina; **Validation**: RP Theart, F Levet, P Hernández-Herrera, LR de la Ballina; **Visualisation**: RP Theart, F Levet, P Hernández-Herrera, LR de la Ballina; **Figure preparation**: LR de la Ballina; **Writing – original draft**: RP Theart, F Levet, LR de la Ballina; **Writing – review & editing**: RP Theart, F Levet, P Hernández-Herrera, C Leterrier, LR de la Ballina.

## DATA AVAILABILITY

Raw, processed and annotated images referred to in this work have been deposited in Zenodo and are publicly available at https://doi.org/10.5281/zenodo.17792816^6, 7^ FIJI plugins used as part of the analyses can be found at https://github.com/kjamartens/thunderstorm for ThunderSTORM and at https://github.com/flevet/WaveJ/releases/tag/1.0.0 for WaveJ.

## Notes

https://doi.org/10.5281/zenodo.17792816

### Competing Interest Statement

The authors have declared no competing interest.

## REFERENCES

1. Schmied, C., Nelson, M. S., Avilov, S., Bakker, G. J., Bertocchi, C., Bischof, J., Boehm, U., Brocher, J., Carvalho, M. T., Chiritescu, C., Christopher, J., Cimini, B. A., Conde-Sousa, E., Ebner, M., Ecker, R., Eliceiri, K., Fernandez-Rodriguez, J., Gaudreault, N., Gelman, L., Grunwald, D., Gu, T., Halidi, N., Hammer, M., Hartley, M., Held, M., Jug, F., Kapoor, V., Koksoy, A. A., Lacoste, J., Le Devedec, S., Le Guyader, S., Liu, P., Martins, G. G., Mathur, A., Miura, K., Montero Llopis, P., Nitschke, R., North, A., Parslow, A. C., Payne-Dwyer, A., Plantard, L., Ali, R., Schroth-Diez, B., Schutz, L., Scott, R. T., Seitz, A., Selchow, O., Sharma, V. P., Spitaler, M., Srinivasan, S., Strambio-De-Castillia, C., Taatjes, D., Tischer, C. & Jambor, H. K. (2024) Community-developed checklists for publishing images and image analyses. Nat Methods, 21, 170–181.

2. Boehm, U., Nelson, G., Brown, C. M., Bagley, S., Bajcsy, P., Bischof, J., Dauphin, A., Dobbie, I. M., Eriksson, J. E., Faklaris, O., Fernandez-Rodriguez, J., Ferrand, A., Gelman, L., Gheisari, A., Hartmann, H., Kukat, C., Laude, A., Mitkovski, M., Munck, S., North, A. J., Rasse, T. M., Resch-Genger, U., Schuetz, L. C., Seitz, A., Strambio-De-Castillia, C., Swedlow, J. R. & Nitschke, R. (2022) Author Correction: QUAREP-LiMi: a community endeavor to advance quality assessment and reproducibility in light microscopy. Nat Methods, 19, 256.

3. Wilkinson, M. D., Dumontier, M., Aalbersberg, I. J., Appleton, G., Axton, M., Baak, A., Blomberg, N., Boiten, J. W., da Silva Santos, L. B., Bourne, P. E., Bouwman, J., Brookes, A. J., Clark, T., Crosas, M., Dillo, I., Dumon, O., Edmunds, S., Evelo, C. T., Finkers, R., Gonzalez-Beltran, A., Gray, A. J., Groth, P., Goble, C., Grethe, J. S., Heringa, J., t Hoen, P. A., Hooft, R., Kuhn, T., Kok, R., Kok, J., Lusher, S. J., Martone, M. E., Mons, A., Packer, A. L., Persson, B., Rocca-Serra, P., Roos, M., van Schaik, R., Sansone, S. A., Schultes, E., Sengstag, T., Slater, T., Strawn, G., Swertz, M. A., Thompson, M., van der Lei, J., van Mulligen, E., Velterop, J., Waagmeester, A., Wittenburg, P., Wolstencroft, K., Zhao, J. & Mons, B. (2016) The FAIR Guiding Principles for scientific data management and stewardship. Sci Data, 3, 160018.

4. Bingham, D., Jakobs, C. E., Wernert, F., Boroni-Rueda, F., Jullien, N., Schentarra, E. M., Friedl, K., Da Costa Moura, J., van Bommel, D. M., Caillol, G., Ogawa, Y., Papandreou, M. J. & Leterrier, C. (2023) Presynapses contain distinct actin nanostructures. J Cell Biol, 222.

5. Burry, R. W. (1980) Formation of apparent presynaptic elements in response to poly-basic compounds. Brain Res, 184, 85–98.

6. OpenAIRE, E. O. F. N. R. a. (2013) Zenodo.

7. Theart, R., Levet, F., Hernandez-Herrera, Leterrier, C., & de la Ballina, L. R. (2025) Sample image data and annotations associated to Presynaptic Actin Nanostructures: A Reproducibility Case Study (v1.0).

8. Leterrier, C., Potier, J., Caillol, G., Debarnot, C., Rueda Boroni, F. & Dargent, B. (2015) Nanoscale Architecture of the Axon Initial Segment Reveals an Organized and Robust Scaffold. Cell Rep, 13, 2781–2793.

9. Ovesny, M., Krizek, P., Borkovec, J., Svindrych, Z. & Hagen, G. M. (2014) ThunderSTORM: a comprehensive ImageJ plug-in for PALM and STORM data analysis and super-resolution imaging. Bioinformatics, 30, 2389–2390.

10. Schindelin, J., Arganda-Carreras, I., Frise, E., Kaynig, V., Longair, M., Pietzsch, T., Preibisch, S., Rueden, C., Saalfeld, S., Schmid, B., Tinevez, J. Y., White, D. J., Hartenstein, V., Eliceiri, K., Tomancak, P. & Cardona, A. (2012) Fiji: an open-source platform for biological-image analysis. Nat Methods, 9, 676–682.

11. Rueden, C. T., Schindelin, J., Hiner, M. C., DeZonia, B. E., Walter, A. E., Arena, E. T. & Eliceiri, K. W. (2017) ImageJ2: ImageJ for the next generation of scientific image data. BMC Bioinformatics, 18, 529.

12. Levet, F., Tonnesen, J., Nagerl, U. V. & Sibarita, J. B. (2020) SpineJ: A software tool for quantitative analysis of nanoscale spine morphology. Methods, 174, 49–55.

13. Holschneider, M., Kronland-Martinet, R., Morlet, J. & Tchamitchian, P. (1990) A Real-Time Algorithm for Signal Analysis with the Help of the Wavelet Transform. Springer Berlin Heidelberg, Berlin, Heidelberg.

14. Wigbers, M. C., Tan, T. H., Brauns, F., Liu, J., Swartz, S. Z., Frey, E. & Fakhri, N. (2021) A hierarchy of protein patterns robustly decodes cell shape information. Nature Physics, 17, 578–584.

15. Sitarska, E., Almeida, S. D., Beckwith, M. S., Stopp, J., Czuchnowski, J., Siggel, M., Roessner, R., Tschanz, A., Ejsing, C., Schwab, Y., Kosinski, J., Sixt, M., Kreshuk, A., Erzberger, A. & Diz-Munoz, A. (2023) Sensing their plasma membrane curvature allows migrating cells to circumvent obstacles. Nat Commun, 14, 5644.

16. Fernholz, M. H. P., Guggiana Nilo, D. A., Bonhoeffer, T. & Kist, A. M. (2024) DeepD3, an open framework for automated quantification of dendritic spines. PLOS Computational Biology, 20, e1011774.

